# Prevalence, distribution, and mechanisms of genetic dominance in the yeast mating pathway

**DOI:** 10.64898/2025.12.16.694484

**Authors:** M. Grace Allen, Artemiza Martínez, Gregory I. Lang

**Affiliations:** Department of Biological Sciences, Lehigh University, Bethlehem PA, USA 18015

**Keywords:** Genetic Dominance, Mutation Rate, *Saccharomyces cerevisiae*, Mating Pathway

## Abstract

The proportion of mutations that are dominant is a fundamental genetic parameter affecting the rate of adaptation, the efficacy of selection, and the maintenance of variation in populations. Yet, estimates for this parameter vary greatly. Here we directly quantify the rates and genetic targets of dominant and recessive mutations in the yeast mating pathway by performing parallel genetic screens in haploid (*MAT***a**) and diploid (*MAT***a**/**a**) yeast. We find that ∼1% of alpha-factor resistant mutations are dominant. We sequenced 95 alpha-factor resistant mutants to determine the genetic architecture of dominant and recessive mutations. We find recessive mutations throughout the mating pathway; however, dominant mutations are concentrated in three genes: *STE4, GPA1*, and *CLN3*. Finally, we investigate the mechanism of genetic dominance in each of these genes. This work sheds light on the prevalence and molecular basis of genetic dominance in a model eukaryotic signal transduction pathway.

## INTRODUCTION

Most loss-of-function mutations are recessive. A systematic analysis in yeast found that only ∼3% of gene deletions show a measurable phenotype when heterozygous (Deutschbauer *et al*. 2005). Dominant mutations, therefore, must act through specific mechanisms of action, for example constitutive activation of a protein or disruption of a protein complex. Classic mutagenesis studies suggest that less than 10% of spontaneous mutations are dominant (Wilkie 1994). However, more recent estimates from Deep Mutational Scanning studies suggest that the fraction of dominant mutations may be as high as a third of all possible mutations (Padhy *et al*. 2023, Flynn *et al*. 2024). Discrepancies in these estimates could be due to specifics of the gene or phenotype under study or the criteria used to assess dominance. One approach to determine the proportion of mutations that are genetically dominant is to measure mutation rates in isogenic haploids and diploids, ideally in a way that samples many genetic targets using the same selection.

The yeast mating pathway is one of the best studied signal-transduction pathways (Herskowitz 1995, Alvaro and Thorner 2016). Signaling is initiated by the mating pheromone (α-factor or **a**-factor) which binds to the cognate receptor (Ste2 or Ste3 for *MAT***a** and *MAT*α cells, respectively). Pheromone binding activates a heterotrimeric G protein, which signals through a MAPK cascade, resulting in both a cell-cycle arrest and a transcriptional response (Bardwell 2005). Mutations that disrupt signaling can therefore be recovered as alpha-factor resistant colonies, providing a simple selection on pathway function. While genetic screens have been instrumental in identifying the components of the yeast mating pathway (Mackay and Manney 1974a, Mackay and Manney 1974b, Manney and Woods 1976, Hartwell 1980), all direct screens for mating mutants have been performed with haploid yeast, as true (*MAT***a**/α) diploids do not have an active mating pathway. Therefore, classic genetic screens for mating defects may have missed rare dominant mutations.

Here, we perform genetic screens to isolate alpha-factor resistance mutants from both isogenic haploid (*MAT***a**) and diploid (*MAT***a/a**) yeast strain to estimate the fraction of resistance mutations that are dominant and to identify the genes and mechanisms that generate dominance in this selection. We characterized 95 alpha-factor resistant mutants by phenotyping and whole-genome sequencing to identify the underlying causative mutations. We find that ∼1% of alpha-factor resistant mutations are dominant. Unlike recessive alpha-factor resistance mutations that are distributed throughout the pathway, dominant mutations are concentrated in three genes: *STE4, GPA1*, and *CLN3*.

## MATERIALS AND METHODS

### Yeast Strains and Plasmids

Strains yGIL432 (haploid *MAT***a**) and yGIL1132 (diploid *MAT***a**/**a**) are derived from the W303 background. yGIL432 has the genotype *ura3*Δ::p*FUS1*::yeVenus *ade2-1, his3-11,15, leu2-3,112, trp1-1, CAN1, bar1*Δ::*ADE2, hml*Δ::*LEU2 GPA1*::NatMX. yGIL1132 is a homozygous diploid derivative of yGIL432. The only exception is at the *URA3* locus, which is heterozygous: *ura*3/*ura3*Δ::p*FUS1*::yeVenus. We assayed mating and **a**-factor production using mating-type testers yGIL355 (*MAT*α, *met4*Δ::KanMX) and yGIL617 (*MAT*α, *sst2*), respectively. We sporulated diploid *MAT***a**/**a** by transforming with the plasmid pGIL069, a *URA3* marked CEN/ARS plasmid with the *MAT*α mating-type information. We constructed *MAT***a**/**a** diploid derivatives of *MAT***a** alpha-factor resistant clones by first complementing the sterile mutation using the appropriate plasmid from the MoBY ORF plasmid collection (Ho *et al*. 2009). We then crossed this strain to yGIL2345, a *MAT*α version of yGIL432 which has KanMX in place of NatMX at the *GPA1* locus and carrying pGIL088, a *URA3* marked plasmid with a galactose-inducible *HO* and the *SpHIS5* gene under control of the *STE2* promoter. We selected for diploids on YPD+G418+ClonNat. Following mating-type switching, we selected *MAT***a**/**a** clones on SC-his plates. Finally, we eliminated the plasmids by counter-selecting on 5FOA.

### Measuring Mutation Rates

We measured mutations rates using Luria-Delbrück Fluctuation Assays as described previously (Lang and Murray 2008, Lang 2018) with minor adjustments. We grew strains in low-glucose (0.2%) YPD medium using 10 μl or 100 μl culture volumes for haploid and diploids, respectively. After one day of growth, we brought the cultures up to 100 μl, if necessary, and spot-plated onto YPD medium containing alpha-factor (10 μg/ml). For each assay we initiated 96 replicate cultures, 24 of which were used to estimate the number of cells per culture and 72 of which were plated to determine the distribution of the number of alpha-factor resistant cells per culture. We analyzed the data from Fluctuation Assays using the Ma-Sandri-Sarkar maximum likelihood method in which the data are fit to a model of the Luria-Delbrück distribution based upon a single parameter *m*, the expected number of mutation events per culture (Sarkar *et al*. 1992). We determined the cell number by hemocytometer and calculated mutation rate using the equation *μ* = *m/N*, where *N* is the average number of cells per culture (approximately equal to the number of cell divisions per culture since the initial inoculum is much smaller than *N*). We assigned 95% confidence intervals on *m* and *μ* using equations 24 and 25 from (Rosche and Foster 2000).

### Whole-Genome Sequencing

We sequenced alpha-factor resistant clones as well as alpha-factor resistant and sensitive pools using 150 nt x 150 nt paired-end sequencing on an Illumina HiSeq 2500 at the Sequencing Core Facility within the Lewis-Sigler Institute for Integrative Genomics at Princeton University. Raw reads were trimmed using trimmomatic/0.36 (Bolger *et al*. 2014) using PE - phred33 parameter. Each sample was aligned to the complete and annotated W303 genome (Matheson *et al*. 2017) using BWA-MEM, v.0.7.15 (Li and Durbin 2009). Variants were identified using FreeBayes/1.1.0 (Garrison and Marth 2012), using default parameters. Variants common to all samples were filtered using the VCFtools/0.1.15 vcf-isec function (v.0.1.12b). Individual VCF files were annotated using SnpEff/5.0 using -formatEff flag (Cingolani *et al*. 2012). We verified the variant calling by viewing individual mutations using Integrated Genome Viewer (Broad Institute). Copy Number Variants (CNVs) were identified using ControlFREEC version 11.5 (Boeva *et al*. 2012). All the text outputs were merged into one data frame to remove all the common calls. We filtered CNV calls by excluding variants that were called more than 10 times, were less than 5 kb, or were located within 5 kb of a telomere or centromere. CNVs and aneuploidies were corroborated by visual inspection of chromosome coverage plots created in R. Briefly, we used samtools-depth to calculate per-site depth from the sorted-bam files. We divided the median chromosome coverage by the median genome-wide coverage using non-overlapping 1000 bp window size and 500 nt step size. The total coverage of the genome was normalized to 1 or 2 for haploid or diploid clones, respectively.

### Phenotypic Assays

Strains yGIL432 and yGIL1132 contain the yEVenus reporter under the control of the *FUS1* promoter downstream of the yeast mating pathway. Using flow cytometry, we assayed basal expression or induced expression following 4 h in 10 μg/ml alpha factor. We assayed **a**-factor production by the ability to produce a halo on a lawn of the halo tester strain yGIL617. We scored alpha-factor hypersensitivity if a strain failed to show robust growth in the presence of pheromone produced by the halo-tester lawn. We assessed mating ability by scoring growth on minimal medium following mating between yGIL432 or yGIL1132 and mating tester strain, yGIL335.

## RESULTS & DISCUSSION

### One percent of the mutations that interfere with the mating pathway are dominant

To determine the relative abundance of dominant and recessive mutations that affect the yeast mating pathway, we performed fluctuation assays to measure the rate of alpha-factor resistance mutations in haploid *MAT***a** and diploid *MAT***a**/**a** backgrounds (Table 1). We find that the mutation rate to alpha-factor resistance in our haploid strain is 2.28 x 10^-6^ per genome per generation (1.75 x 10^-6^, 2.86 x 10^-6^, 95% CI). This assay was performed in an *hml*αττ strain to prevent expression of the *MAT*α information due to mating-type switching or loss of silencing.

**Table 1:** Mutation rates to alpha-factor resistance.

| Strain | Ploidy | Relevant Genotype | Mutation Rate | 95% Conf. Int. | Fold relative to yGIL432 |
| --- | --- | --- | --- | --- | --- |
| yGIL104 | Haploid | *MATa HMLα* | $5.86 \times 10^{-6}$ | $5.46 \times 10^{-6} - 6.28 \times 10^{-6}$ | 2.57 |
| yGIL432 | Haploid | *MATa hml$\alpha\Delta$* | $2.28 \times 10^{-6}$ | $1.75 \times 10^{-6} - 2.86 \times 10^{-6}$ | 1.00 |
| yGIL1132 | Diploid | *MATa/a hml$\alpha\Delta$/hml$\alpha\Delta$* | $3.10 \times 10^{-8}$ | $1.31 \times 10^{-8} - 5.41 \times 10^{-8}$ | 0.01 |

Previous measurements of the rate of alpha-factor resistance in an *HML*α strain are two-fold higher (Table 1), consistent with estimates of mating type switching at a rate of ∼10^-6^ per genome per generation (Hicks and Herskowitz 1976, Strathern and Herskowitz 1979). We find that the mutation rate to alpha-factor resistance in *MAT***a**/**a** diploids is 3.10 x 10^-8^ per genome per generation (1.31 x 10^-8^, 5.41 x 10^-8^, 95% CI), two orders of magnitude lower than the rate in haploid cells. This suggests that ∼1% of mutations that confer alpha-factor resistance are dominant, much lower than the estimate of ∼30% from systematic studies of deep mutational scanning (Padhy *et al*. 2023, Flynn *et al*. 2024) and slightly lower than the estimate of <10% from surveys of spontaneous mutations (Wilkie 1994).

### Most alpha-factor resistant diploid clones harbor a single dominant alpha-factor resistance mutation

We selected 47 diploid and 48 haploid alpha-factor resistant clones for further study. We performed tetrad dissections on twenty of the diploid clones to determine the segregation pattern of alpha-factor resistance (Table 2). Three clones (Diploid A07, A08, and C11) showed mixed segregation patterns for spore viability and alpha-factor resistance. These clones were later found to harbor mutations in *CLN3* (Table 2) which interacts with *IME1*, the master regulator of meiosis (Colomina *et al*. 1999). One clone (Diploid D09) produced only alpha-factor resistant spores, consistent with a homozygous mutation, whereas two clones (Diploid A12 and G09) produced only alpha-factor sensitive spores. To determine whether haploid clones carry recessive alpha-factor resistance mutations, we backcrossed four haploid clones (Haploid B01, D02, D03, and G03) to a wild-type strain and gene-converted the *MAT* locus to assay alpha-factor sensitivity (Figure S1). All four backcrossed haploid clones are sensitive to alpha-factor. Combined, these experiments are consistent with single dominant alpha-factor resistant mutations arising in the *MAT***a**/**a** diploid background and recessive alpha-factor resistant mutations in the haploid *MAT***a** background.

**Table 2:** Segregation patterns for alpha-factor resistant diploid clones.

| Clone | Mutations | Causative Mutation | Segregation Patterns |  |
| --- | --- | --- | --- | --- |
| | | | Viability | αF Resistance |
| Diploid A07 | 1 | *CLN3* | 4:0 | Mixed (2:2, 3:1) |
| Diploid A08 | 2 | *CLN3* | Mixed (4:0, 3:1, 2:2) | Mixed (20/27 αFR) |
| Diploid A12 | 0 |  | 4:0 | 0:4 |
| Diploid B07 | 1 | *GPA1* | 4:0 | 2:2 |
| Diploid B08 | 2 |  | 4:0 | 2:2 |
| Diploid B09 | 2 | *GPA1* | 4:0 | 2:2 |
| Diploid B10 | 2 | *GPA1* | 4:0 | 2:2 |
| Diploid B12 | 1 |  | 4:0 | 2:2 |
| Diploid C08 | 1 | *GPA1* | 4:0 | 2:2 |
| Diploid C09 | 3 | *STE4* | 4:0 | 2:2 |
| Diploid C10 | 1 | *STE4* | 4:0 | 2:2 |
| Diploid C11 | 1 | *CLN3* | 4:0 | Mixed (3:1, 2:2) |
| Diploid D08 | 2 | *GPA1* | 2:2 | 2:2 |
| Diploid D09 | 3 | *STE11* | 2:2 | 2:0 |
| Diploid D10 | 2 | *GPA1* | 4:0 | 2:2 |
| Diploid D12 | 2 |  | 4:0 | 2:2 |
| Diploid E10 | 2 | *STE4* | 4:0 | 2:2 |
| Diploid F12 | 0 |  | 2:2 | 2:2 |
| Diploid G09 | 2 |  | 4:0 | 0:4 |
| Diploid G11 | 3 |  | 2:2 | 2:2 |

### Recessive and dominant mutations affect different parts of the mating pathway

We sequenced all 47 diploid and 48 haploid alpha-factor resistant clones, as well as pooled sensitive and resistant spores from tetrad dissections of diploid clones C08, C09, C10, D08, D09 and D10. We find a similar number of mutations in the haploid and diploid alpha-factor resistant clones:1.50 mutations (range of 0 to 5) for the haploid clones and 1.45 mutations (range of 0 to 3) for the diploid clones. For 45 of the 48 haploid clones and 33 of the 47 diploid clones, we could unambiguously identify the causative alpha-factor resistance mutation. In all cases but one, these were non-synonymous changes to proteins associated with the yeast mating pathway or cell-cycle progression. The only exception is a mutation upstream of the *STE4* gene in the haploid clone G05; this mutation appears to create an out-of-frame start codon 50 bp upstream of the annotated start site and is therefore predicted to behave as a null mutation. Among diploid clones, 26 of the 33 candidate causative mutations are heterozygous, consistent with the results from the tetrad dissections which showed 2:2 (or close to 2:2) segregation of alpha-factor resistance for 17 of the 20 diploid clones (Table 2).

Excluding the homozygous mutations in the diploid clones, alpha-factor resistance mutations in haploids and diploids affect distinct parts of the yeast mating pathway (Figure 1). Recessive mutations (those arising in haploids or present as homozygous in the diploid) are found throughout the pathway, with the highest enrichment in the MAPK cascade (*STE11, STE7*, and *STE5*) and the transcription factor (*STE12*). In contrast, dominant mutations are found concentrated in the Gα and Gβ subunits (encoded by *GPA1* and *STE4*, respectively) as well as the cyclin gene, *CLN3*.

**Figure 1.**
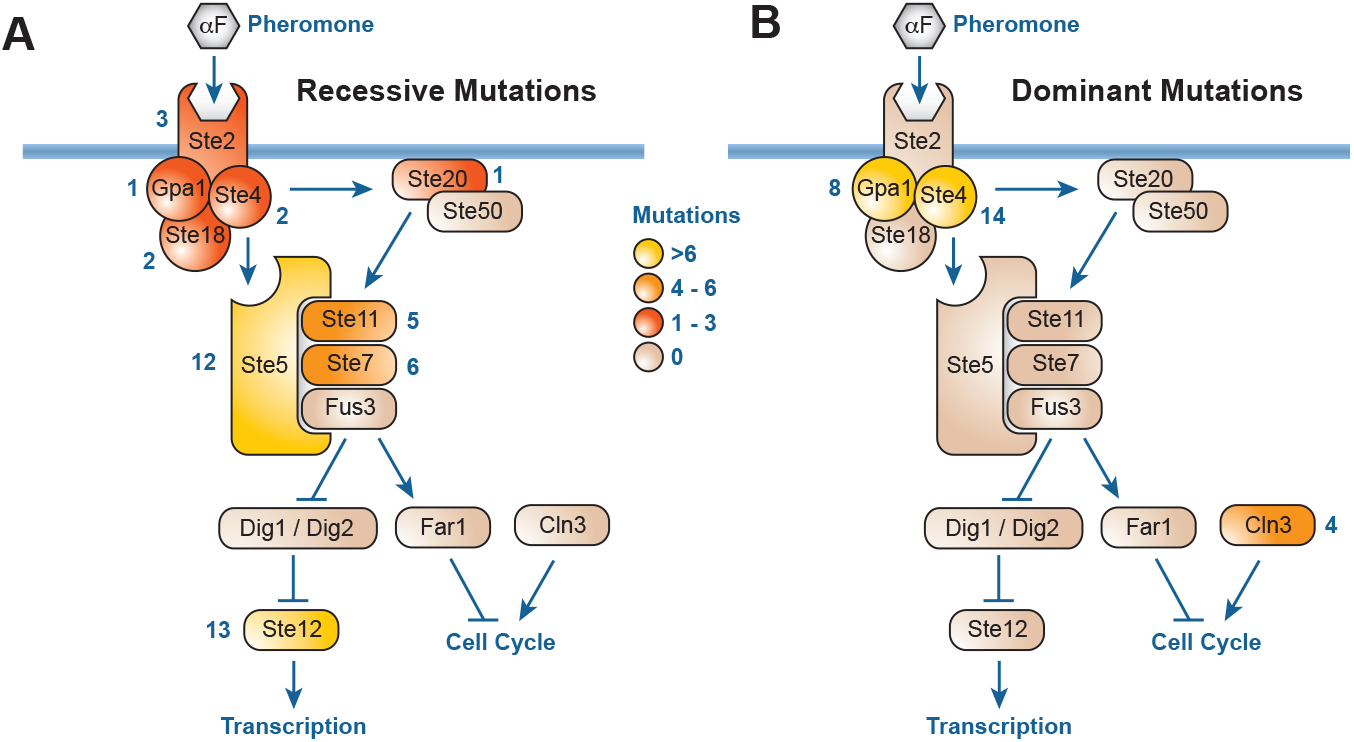
Recessive and dominant mutations affect different parts of the mating pathway. Schematic of the yeast mating pathway showing the number of times each gene was identified as the putative alpha-factor resistant mutation in the *MAT***a** (recessive) and *MAT***a**/**a** (dominant) genetic screens, respectively. **(A)** Recessive mutations are found throughout the mating pathway. **(B)** Dominant mutations are found only in the Gα and Gβ subunits (encoded by *GPA1* and *STE4*, respectively) as well as the cyclin gene, *CLN3*. Seven mutations from the *MAT***a**/**a** screen that underwent loss-of-heterozygosity are not shown.

### Some diploid clones contain additional recessive mutations

In diploid clones, we observed seven cases of loss of heterozygosity (LOH) – *STE2* in A09, B11, G10, H11; *STE5* in H08; and *STE11* in D09 and F07 – and at least four clones (A09, B11, G10, G11) that harbor one or more aneuploidies (Supplemental Dataset 1 and Figure S2). The number of homozygous mutations is somewhat surprising given that only ∼20 generations of growth occurred prior to plating on selective medium and homozygous mutations require a two-step process (mutation followed by LOH). Two homozygous mutations occurred in *STE11*, which lies distal to the rDNA repeat on Chromosome XII, a known hotspot for mitotic recombination (Marad *et al*. 2018). Five homozygous mutations in *STE2* on Chromosome VI; these were accompanied by loss of Chromosome VI and Chromosome I (Figure S2). Given that about a quarter (12/47) of the alpha-factor resistance mutations in the diploids are recessive, the mutation rate to dominant resistance (2.31 x 10^-8^) is slightly less than the mutation rate to alpha-factor resistance in diploids (3.10 x 10^-8^, Table 1).

Homozygous mutations and aneuploidies also explain some of the unexpected segregation patterns (Table 2). Most clones showed were consistent with a single dominant alpha-factor resistant mutation: four-spore viable tetrads segregating 2:2 alpha-factor resistance. However, four of the twenty clones showed 2:2 segregation for viability, indicating that, besides the resistant mutation, these clones also harbor a recessive lethal mutation, which could arise from a point mutation or the loss of an entire chromosome. For diploid clone D09, we observe 2:2 segregation of viability and 2:0 segregation of alpha-factor resistance. This clone has a normal karyotype, a homozygous *STE11* mutation, and heterozygous missense mutation in *RIB4* and *UBC7* (Supplemental Dataset 1). Neither *RIB4* nor *UBC7* are annotated as essential genes in the Saccharomyces Genome Database. To identify the recessive lethal mutation in clone D09, we sequenced the pooled viable spores. We find that, while *UBC7* is present in the viable spores, the *RIB4* mutation is absent, indicating that *RIB4* is essential in our strain background.

### Dominant and recessive alpha-factor resistance mutations fall into distinct phenotypic clusters

We measured phenotypes associated with mating ability for all haploid and diploid alpha-factor resistant mutants, including basal and induced expression using a p*FUS1*-yEVenus reporter, **a**-factor production, and mating ability. We also assessed pheromone sensitivity in two ways: by replating on YPD+alpha factor and by spotting on a lawn of *MAT*α cells (Figure 2). Clustering the phenotypes produces expected relationships: basal expression clusters with a-factor production and induced expression clusters with mating ability. Interestingly, 22 of the 47 diploid clones initially isolated on YPD+alpha factor arrested when retested, despite most of them having a confirmed heterozygous mutation in the mating pathway, suggesting reduced penetrance of dominant resistance mutations.

**Figure 2.**
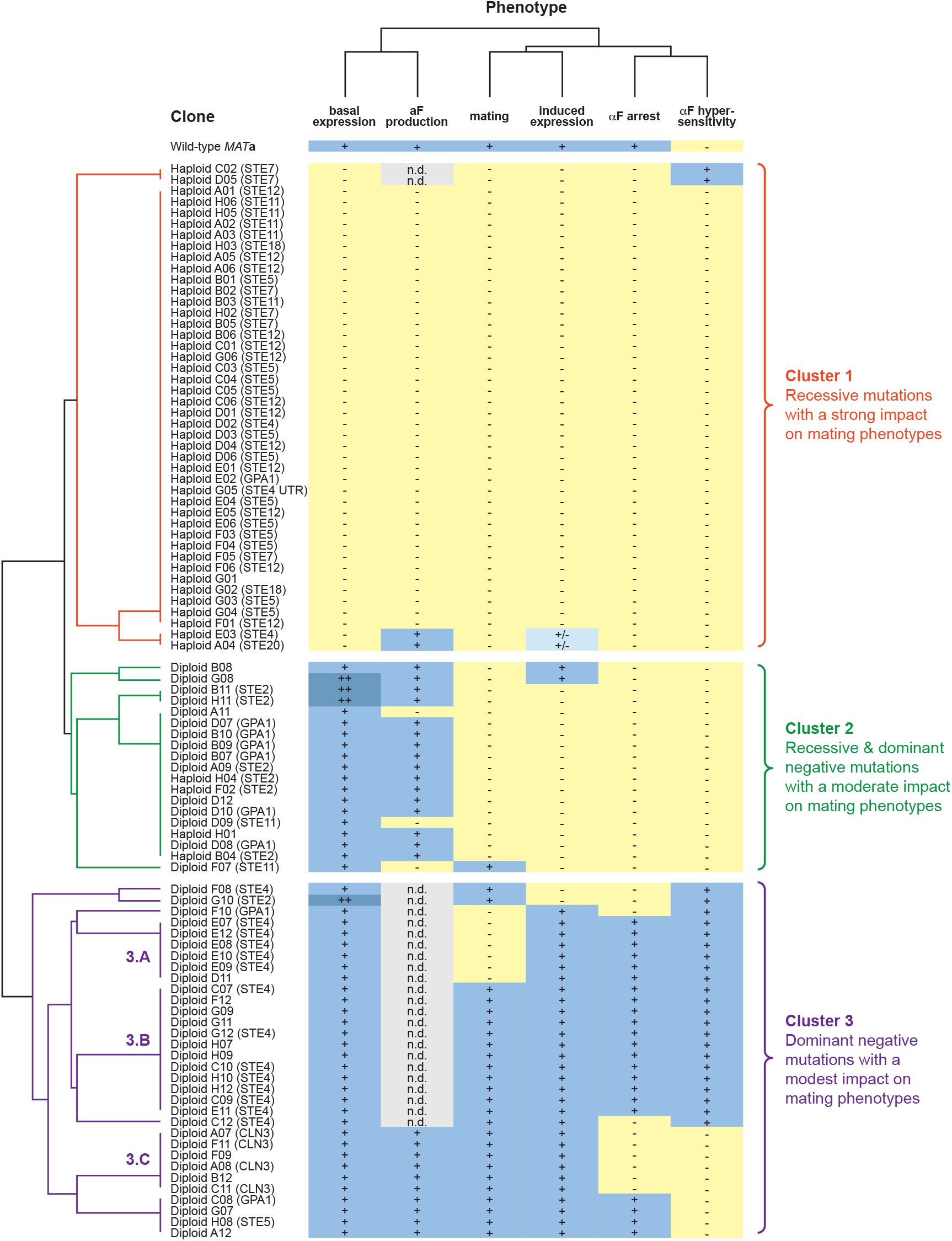
Diverse phenotypes among sterile mutants. We measured six phenotypes associated with mating ability for all haploid and diploid alpha-factor resistant mutants. All phenotypes were binary, with a few exceptions. We were unable to assay **a**-factor production for strains that were hypersensitive to alpha-factor. Several strains displayed higher than WT basal expression (++), and some strains displayed intermediate levels of induced expression (+/-). We clustered the data by both clone and by phenotype.

Resistant clones formed three main clusters. Cluster 1 are haploid clones harboring recessive null mutations in the pathway; these clones showed strong mating defects and lack detectable basal expression or mating ability. Mutations in Cluster 1 occurred throughout the pathway but are particularly enriched in the MAPK pathway genes *STE11* (5/44), *STE7* (6/44), *STE5* (12/44) and the transcription factor gene *STE12* (13/44). Cluster 2 includes both haploid and diploid clones with intermediate phenotypes: clones show basal expression through pathway but lack either induced expression or mating ability. Mutations in Cluster 2 tend to fall in the alpha-factor receptor gene *STE2* (6/19), or the Gα subunit gene, *GPA1* (6/19). Cluster 3 contains diploid clones with modest effects on mating-related phenotypes; mutations were concentrated in the Gβ subunit gene, *STE4* (14/32), and the G1 cyclin gene, *CLN3* (4/32).

### Dominant mutations in *GPA1* and *CLN3* are gain-of-function

The identities and patterns of mutations in *GPA1* and *CLN3* imply specific molecular mechanisms for dominant alpha-factor resistant mutations. Dominant *GPA1* mutations cluster in two regions. The first region is in the GTP binding domain. Prior work showed that mutations at residues Gly322, Arg327, and Glu364 (the “G-R-E motif”) in Gpa1 (Gα) stabilize the GDP-bound configuration, locking the protein in the inactive state and preventing Ste4/Ste18 (Gβγ) release (Knight *et al*. 2021). Adjacent mutations (Gln323Lys and Asn233Asp) produce a similar phenotype (Wu *et al*. 2004). Consistent with this, we observe three substitutions at Gly322 (Gly322Val, Gly322Arg, Gly322Glu) and one at Glu364 (Glu364Lys; Figure 3). While we do not find mutations at Arg327, we do observe a mutation at an adjacent, and well-conserved, arginine residue (Arg324Leu).

**Figure 3.**
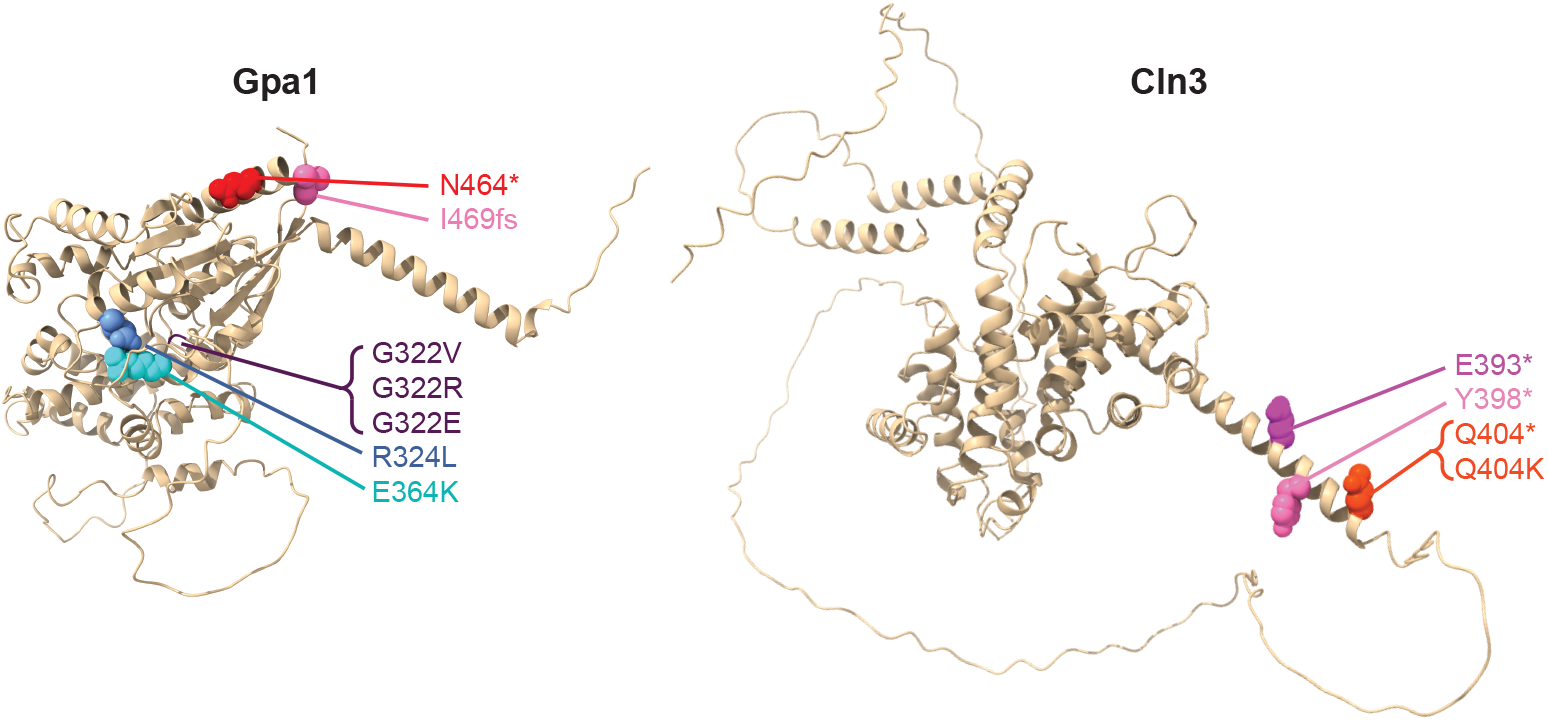
Dominant mutations in Gpa1 and Cln3. The identity and patterns of mutations in *GPA1* and *CLN3* imply specific molecular mechanisms for dominant alpha-factor resistant mutations. Dominant mutations in Gpa1 cluster in two regions: the GTP binding domain and the C-terminal domain that interacts with the Ste2 receptor. Dominant mutations in Cln3 arise in (or truncate) the C-terminal PEST domain.

A second group of dominant *GPA1* mutations cluster in the C-terminal α5 helix, which is the primary point of contact between Gpa1 and the Ste2 receptor (Velazhahan *et al*. 2021).

Missense mutations in this region modulate the strength of signaling through the mating pathway. Lysine-to-proline substitutions at positions 467 and 468, or a nonsense mutation at position 468, decrease mating ability and induced signaling through the mating pathway (Hirsch *et al*. 1991, Yvert *et al*. 2003). We identify a nonsense mutation at position 464 and a frameshift at position 469 that add six amino acids to the C-terminus of the protein (Figure 3). These are likely gain-of-function mutations that interfere with release of the G protein from the Ste2 receptor, because loss of this interaction (for example by deletion of *GPA1*) would result in constitutive signaling through the pathway and a cell-cycle arrest in the absence of alpha-factor.

Another common target for dominant mutations is the G1 cyclin gene *CLN3*. All four dominant *CLN3* mutations affect the C-terminal PEST domain (Figure 3). Mutations in this region are known to be hyperactive dominant alleles that shorten G1 and decrease alpha-factor sensitivity. One mutation changes Lys404 to Gln; and the other three remove the PEST domain. Interestingly, one truncation (Q404*) is identical to the original *cln3-1* allele (Nash *et al*. 1988). Loss of the PEST domain produces a more stable, hyperactive cyclin that promotes G1 to S progression (Cross and Blake 1993). Like the dominant *GPA1* mutants, our dominant *CLN3* mutants are not dominant-negative, as null mutations of *CLN3* have the opposite phenotype from what we observe: prolonged G1 phase and sensitivity to alpha-factor (Teufel *et al*. 2019).

### Dominant mutations in *STE4* are dominant-negative

Ste4 is the Gβ subunit of the heterotrimeric G-protein. It is a WD40 domain-containing protein that adopts a canonical seven-blade beta-propeller structure (Figure 4A). Over half of the dominant alpha-factor resistant mutations (14/26) are found in the *STE4* gene. The nine nonsense/frameshift and five missense mutations are distributed throughout the protein (Figure 4B). Dominant missense mutations tend to be in phenotypic Cluster 3.A, whereas nonsense/frameshift tend to be Cluster 3.B (Figure 4C). Both Cluster 3.A and 3.B have basal and induced signaling through the mating pathway; however, Cluster 3.A is unable to mate (Figure 2). In *STE4*, therefore, dominant missense mutations have more severe phenotypes than dominant nonsense/frameshift mutations, a pattern consistent with a dominant-negative mechanism in which a subset of missense mutations interfere with the function of the protein complex.

**Figure 4.**
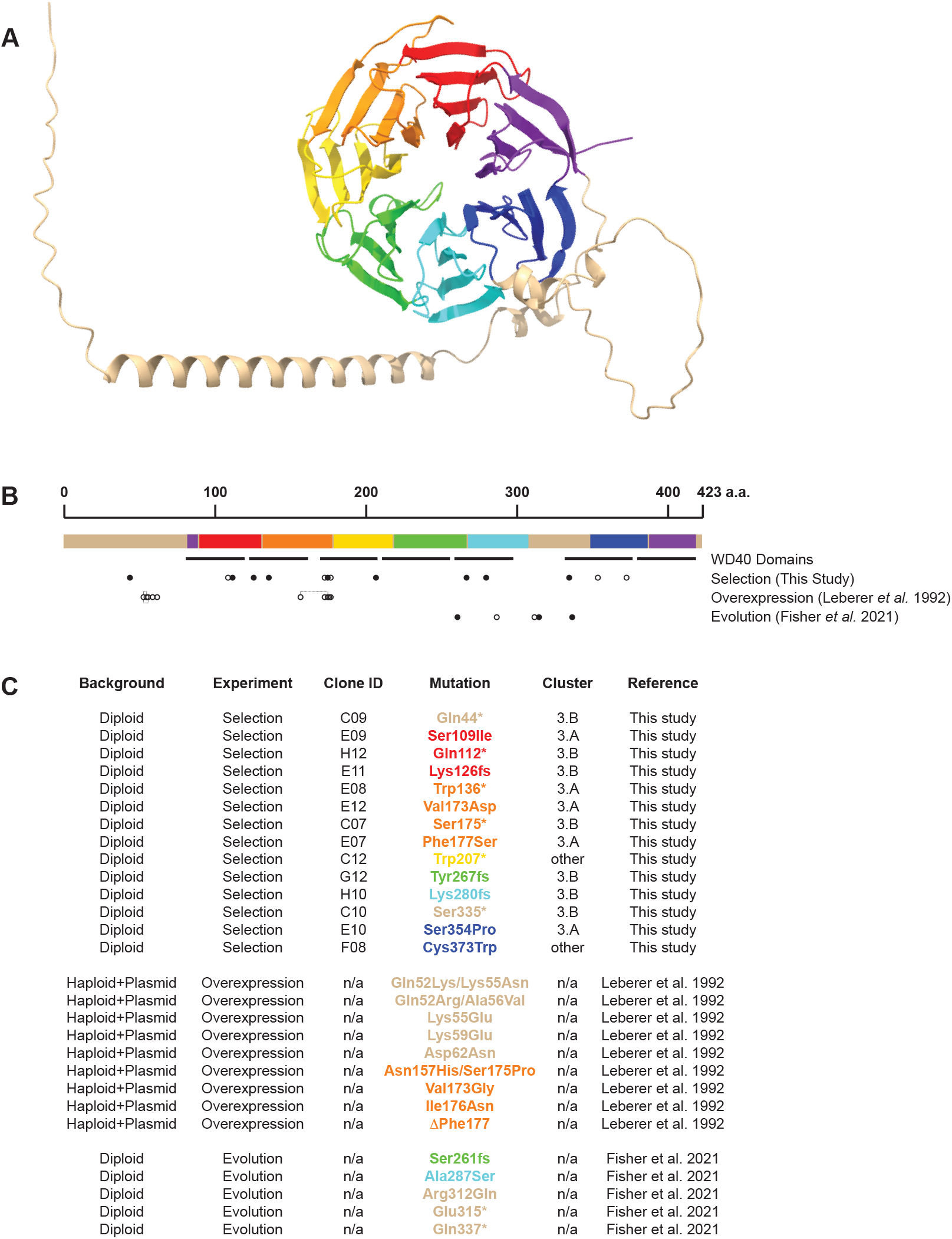
Dominant-negative mutations are distributed throughout the Ste4 protein. **(A)** Ste4 is a WD40 domain-containing protein that adopts a seven-blade beta-propeller structure. The structure shown is the alpha-fold predicted structure with each blade indicated a different color. **(B)** The primary structure of the Ste4 protein indicating the locations of the WD40 domains, and the dominant-negative missense (open circles) and nonsense/frameshift mutations (filled circles) from this study, as well as dominant-negative mutations from an overexpression study (Leberer *et al*. 1992) and from experimental evolution (Fisher *et al*. 2021). **(C)** A list of dominant-negative Ste4 mutations from this study, Leberer *et al*. 1992, and Fisher *et al*. 2021.

An earlier study overexpressed a mutagenized *STE4* library in a haploid strain to screen for dominant-negative mutations affecting alpha-factor resistance (Leberer *et al*. 1992). This study identified nine dominant-negative missense mutations in Ste4, but unlike our results, they observed mutations in only two regions of the protein. Interestingly, we observe mutations at two of the residues found in this prior study: Val173 and Phe177. Both our results and Leberer *et al*. 1992 indicate that dominant-negative mutations in *STE4* have modest effects on other mating-associated phenotypes, including basal and induced expression of downstream reporters and mating ability.

The observation that *STE4* is a large target for alpha-factor resistant mutations is consistent with our previous results from experimental evolution. Mutations that inactivate the mating pathway are common in laboratory evolution experiments and confer a fitness benefit due to elimination of the cost of basal gene expression. *MAT***a**/**a** autodiploids, like haploids, should benefit from mutations that inactivate the mating pathway (Lang *et al*. 2009). However, only *STE4* is identified as a common target of selection across the autodiploid populations (Fisher *et al*. 2018, Fisher *et al*. 2021).

Overall, our results show that for the yeast mating pathway, approximately 1% of spontaneous mutations are dominant. These dominant mutations are clustered in the Gα and the Gβ subunits of the heterotrimeric G-protein (Gpa1 and Ste4, respectively) as well as the G1-cyclin (Cln3). Mutations in *STE4* appear to be dominant-negative, interfering with signaling. Mutations in *GPA1* and *CLN3* appear to be dominant mutations that lock these proteins in a state that either prevents signaling through the pathway or allows for bypass of the pheromone-induced G1 arrest. These results shed light on the prevalence of dominant mutations and the mechanisms of genetic dominance.

## DATA AVAILABILITY

The short-read sequencing data reported in this study have been deposited to the NCBI BioProject database, accession number PRJNA1330868.

## ACKNOWLEDGEMENTS AND FUNDING

We thank members of the Lang Lab for comments on the manuscript. This work was supported by grants from the National Institutes of Health (R35GM149540 and R01GM127420). We are grateful to the NIH for an Administrative Supplement to Support Undergraduate Summer Research (R01GM127420-04S1). Portions of this research were conducted on Lehigh University’s Research Computing infrastructure partially supported by the National Science Foundation (Award 2019035).

## AUTHOR CONTRIBUTIONS

Conceptualization: GIL

Experiments: MGA, AAM, GIL

Analysis: MGA, GIL

Writing: GIL

Editing: GIL, AAM, MGA

Funding Acquisition: GIL

---

**Supplementary Figure 1.**
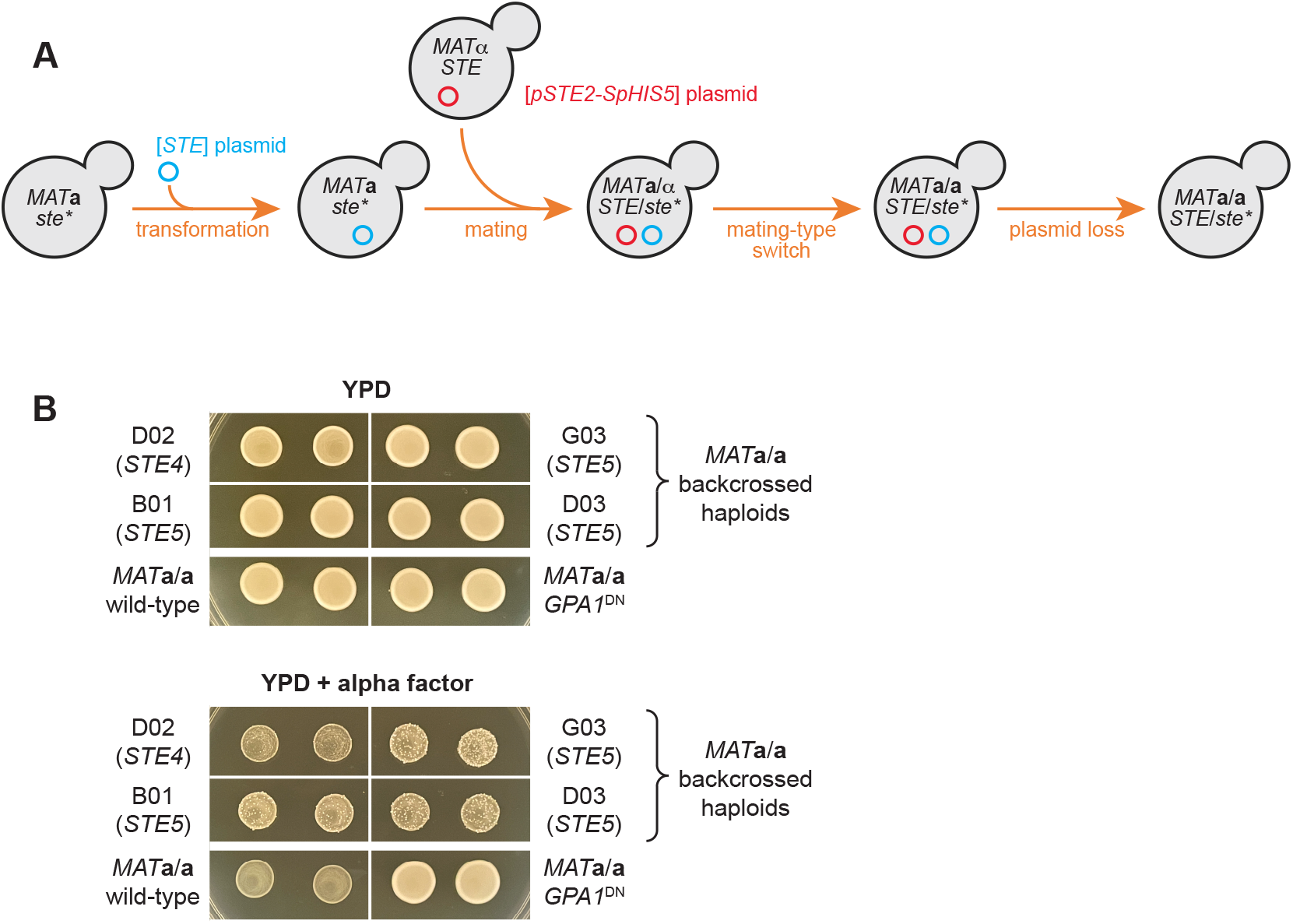
Alpha-factor resistance mutations arising in haploids are recessive. **(A)** We complemented sterile mutations in the alpha-factor resistant clones using the appropriate plasmid from the MoBY ORF plasmid collection (Ho *et al*. 2009) and mated each to a *MAT*α strain containing a plasmid with a galactose-inducible HO and the *SpHIS5* gene under control of the *STE2* promoter. Following mating-type switching, *MAT***a**/**a** clones were selected on SC-his plates, and plasmids were eliminated by plating to 5FOA. **(B)** Growth of strains heterozygous for *ste4* or *ste5* mutations that arose in the haploid background. The *ste4* heterozygous mutations grows slower, consistent with our previous observation that heterozygous *STE4/ste4*Δ mutants are underdominant (Fisher *et al*. 2021). The control clone with a dominant negative allele of *GPA1* grows well on both media; however, the control wild-type *MAT***a**/**a** clone and the four backcrossed haploid clones heterozygous for either *ste4* or *ste5* mutations do not grow on YPD+alpha factor. The breakthrough mutations on backcrossed haploid clones are likely due to loss of heterozygosity, which is expected to occur at a high frequency.

**Supplementary Figure 2.**
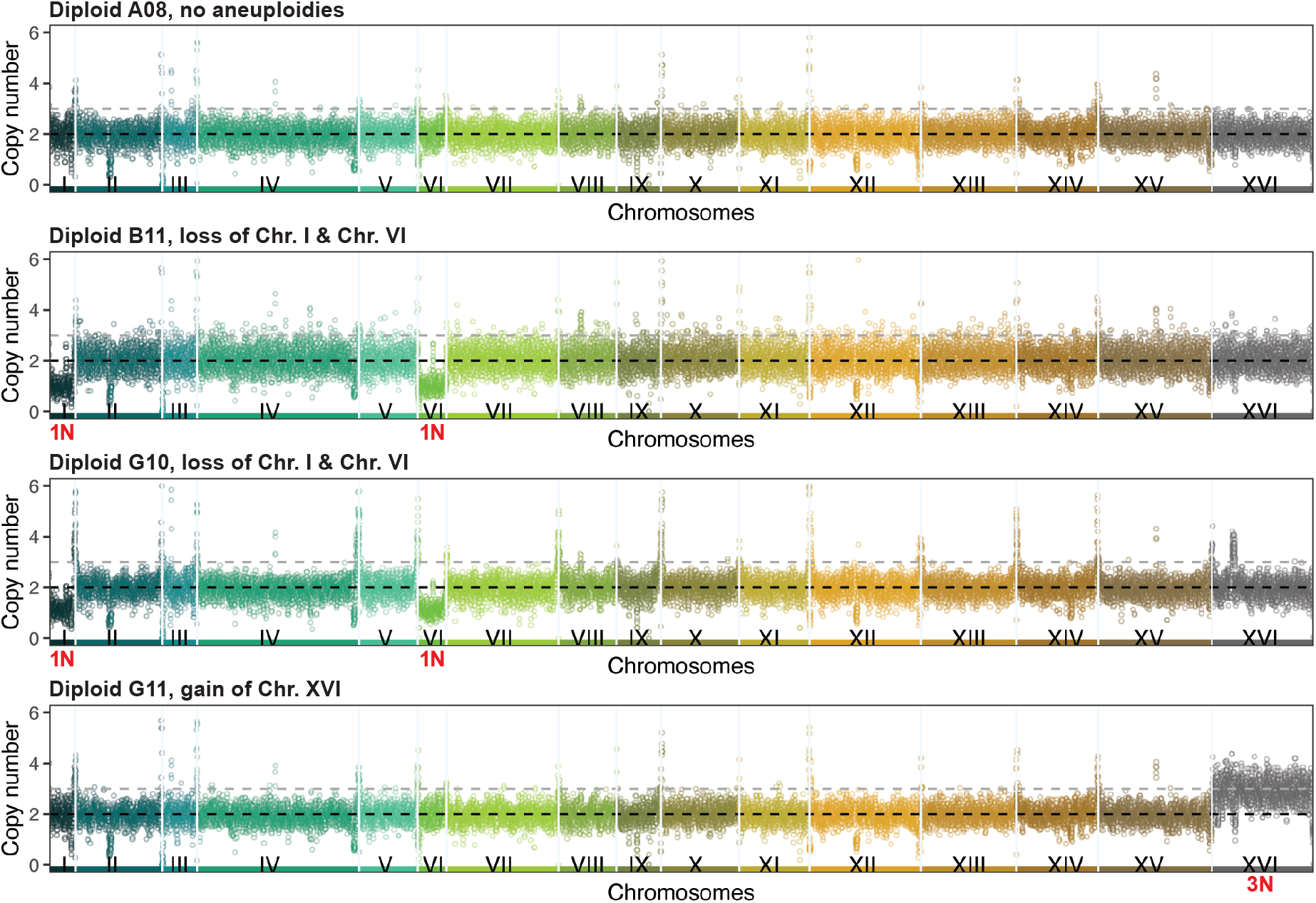
Aneuploidies in alpha-factor resistant *MAT*a/a diploid strains. Coverage across each position on the genome using a sliding 1 kb sliding window with a step of 500 bp. Each chromosome is represented by a different color. The baseline ploidy was normalized to 2N.

